# Vasopressin V2 receptor phosphopeptide reshapes the conformational landscape of arrestin-2 beyond C-tail detachment

**DOI:** 10.64898/2026.11.05.764624

**Authors:** Maria Bzówka, Raphaël Feser, Simone Aureli, Layara Akemi Abiko, Francesco Luigi Gervasio

**Affiliations:** School of Pharmaceutical Sciences, University of Geneva, CH-1206 Geneva, Switzerland; Institute of Pharmaceutical Sciences of Western Switzerland, University of Geneva, CH-1206 Geneva, Switzerland; Swiss Bioinformatics Institute, University of Geneva, CH-1206 Geneva, Switzerland; Neuroscience, Physiology and Pharmacology Department, University College London (UCL), WC1E 6BT London, U.K.; Chemistry Department, University College London (UCL), WC1E 6BT London, U.K.

**Author notes:** These authors contributed equally to this work (Maria Bzówka, Raphaël Feser).

## Abstract

Binding of phosphorylated GPCR tails influences arrestin conformation, but the contribution of this interaction to the interdomain rotation associated with activation remains incompletely characterised. In particular, it remains unclear whether displacement of the arrestin C-tail is sufficient to favor larger interdomain rotations or whether the bound phosphotail exerts an additional effect on the conformational landscape. Here, we used enhanced-sampling simulations to characterise the conformational free-energy landscapes of basal (unbound) and V2Rpp-bound arrestin-2. We find that V2Rpp binding broadened the range of energetically accessible interdomain rotation angles relative to basal arrestin-2. However the pronounced rotation observed in the experimental structures, while accessible, was not strongly favored. Strikingly, C-tail displacement alone does not reproduce this effect, as arrestin-2 retains a strong preference for small interdomain rotation angles in the absence of V2Rpp. Moreover, the C-tail-detached state is energetically unfavorable in the absence of V2Rpp. Together, these findings quantify the effects of V2Rpp binding on the interdomain rotation of arrestin-2 and disentangle the effects of phosphopeptide binding from those of C-tail detachment.

## 1 Introduction

G protein-coupled receptors (GPCRs) are membrane proteins that transduce extracellular stimuli into intracellular signals through interactions with a variety of cytoplasmic partners, including heterotrimeric *G-proteins, G-protein-coupled receptor kinases* (GRKs), and *arrestins* [1, 2, 3, 4, 5, 6, 7]. Among these, arrestins play central roles in GPCR regulation by promoting receptor desensitization and internalization and by modulating downstream signaling responses [8, 9, 10, 11]. Of the four mammalian arrestin proteins, the two non-visual members, arrestin-2 and arrestin-3, also known as *β*-arrestin 1 and *β*-arrestin 2, are broadly expressed and regulate diverse GPCRs. This broad receptor coverage has made questions about arrestin conformation a central topic in GPCR pharmacology [12, 13, 14, 15, 16]. Structurally, arrestin is composed of two globular domains, the *N-domain* and *C-domain*, connected by an interdomain hinge. Together, these domains form the receptor-binding surface and contain several activation-sensitive structural elements, including the *finger, middle*, and *lariat* loops (Fig. 1A) [17]. In the basal state, arrestin adopts an autoinhibited conformation stabilised by a network of intramolecular interactions that couple the two domains. A key component of this network is arrestin’s C-tail, whose *β*20 strand binds the *β*1 element located in the N-domain and contributes to stabilization of the basal architecture [18, 12, 19, 20].

**Figure 1:**
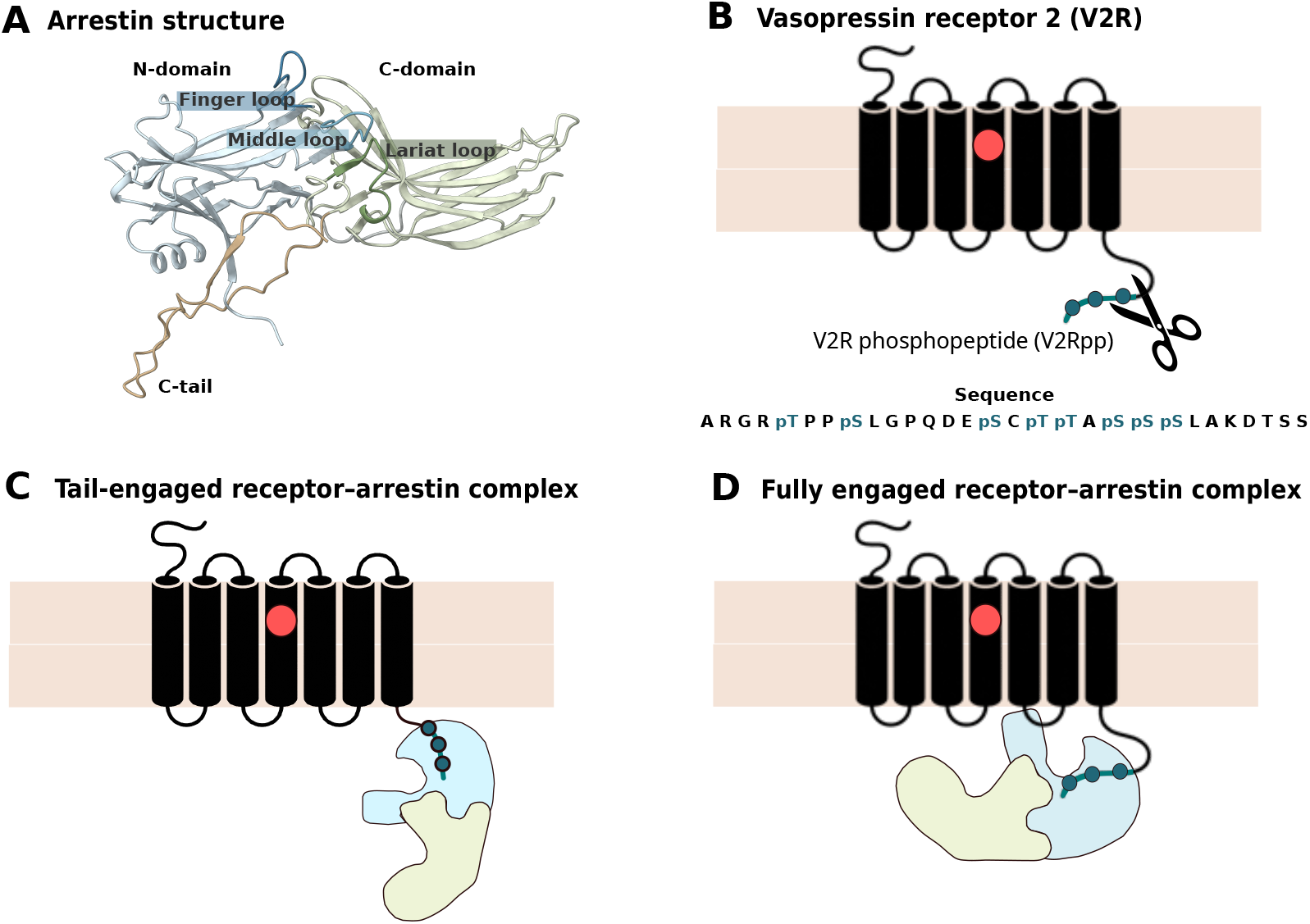
Schematic representation of arrestin-2 structure, V2Rpp, and models of arrestin-2 recruitment. **(A)** Structure of arrestin-2 highlighting the N- and C-domains, C-tail, and key functional loops, i.e. the finger, middle, and lariat loops. **(B)** The V2Rpp sequence with the phosphorylation sites indicated. **(C)** Tail-engaged model of arrestin activation, in which only the phosphotail is involved in arrestin binding. **(D)** Fully-engaged model, in which arrestin is recruited by both phosphotail and receptor core.

Arrestin engagement with GPCRs is promoted primarily by two signals, namely the activated conformation of the GPCR core and the phosphorylated receptor C-terminal tail, hereafter referred to as the phosphotail (Fig. 1). In fully engaged complexes, arrestin contacts both the receptor core and the phosphotail simultaneously. In tail-engaged complexes, by contrast, arrestin binds the phosphotail without stable engagement of the receptor core [21, 22, 14, 16]. Although receptor activation commonly favors arrestin engagement, it is not an absolute prerequisite, as inactive receptor conformations can also support arrestin binding, including core engagement with *β*_1_ adrenergic receptor (*β*_1_AR) [16] and tail engagement with glucagon receptor (GCGR) [23]. One of the most prominent conformational rearrangements associated with arrestin-receptor engagement is interdomain rotation, defined as rotation of the C-domain relative to the N-domain. This rearrangement has been observed in fully engaged receptor–arrestin complexes and is associated with changes in membrane-facing regions and surfaces implicated in downstream-partner interactions, making it an informative structural readout of arrestin activation [18, 24, 25, 26]. Whether comparable interdomain rearrangements are also promoted by tail-only engagement remains less clear. This distinction is important because phosphotail binding and arrestin C-tail displacement are closely coupled. Binding of a phosphotail can displace the autoinhibitory arrestin C-tail from its basal binding site, but it remains unclear whether the resulting conformational changes arise from C-tail detachment itself or from an additional allosteric effect of the bound phosphotail. If C-tail detachment alone were sufficient to drive substantial interdomain rotation, this conformational change would represent a downstream consequence of relief from autoinhibition. Conversely, if phosphotail binding reshapes the interdomain free-energy landscape beyond the effect of C-tail detachment, then the receptor phosphotail would act as an active allosteric regulator of arrestin conformation.

The vasopressin V2 receptor phosphopeptide (V2Rpp) provides a minimal system in which these contributions can be distinguished in the absence of receptor-core engagement (Fig. 1). Tail-engaged receptor–arrestin complexes can support particular endocytic and signaling responses, making their conformational properties relevant beyond an intermediate step toward full engagement [27, 21]. The crystal structures of V2Rpp-bound arrestin-2 show a pronounced interdomain rotation [18, 14]. However, these structures were obtained with Fab30, a conformationally selective antibody fragment that shifts arrestin’s conformational equilibrium toward activated-like states, complicating interpretation of the rotational effect of V2Rpp alone [28, 29]. Nevertheless, Fab-free solution NMR shows that V2Rpp displaces *β*20 and alters arrestin-2 loop conformations and interdomain organization [16]. How V2Rpp binding changes the relative energetic accessibility of interdomain orientations, and, whether C-tail detachment alone accounts for this effect, remain unresolved.

Here, we address these questions using enhanced-sampling molecular dynamics (MD) simulations. We compare the conformational free-energy landscapes of basal and V2Rpp-bound arrestin-2, focusing on interdomain rotation. We then compare these landscapes with that of C-tail-detached arrestin-2 to distinguish the effect of V2Rpp binding from that of C-tail detachment alone. Finally, we quantify the free-energy cost of C-tail detachment in basal arrestin-2 to assess whether the detached state is energetically accessible in the absence of V2Rpp.

## 2 Results

### 2.1. Basal and V2Rpp-bound arrestin-2 retain their characteristic interactions

We first compared the dynamics of the basal and V2Rpp-bound arrestin-2 models using four independent 10 *µs* conventional MD simulations for each system, yielding 40 *µ*s per state and 80 *µ*s in total. This initial analysis tested whether the models retained the structural stability and interaction signatures expected for their respective basal and V2Rpp-engaged states. In basal arrestin-2, the reconstructed proximal C-terminal segment preserved chain continuity around the resolved *β*20 strand, which is part of the autoinhibitory C-tail. In the V2Rpp-bound model, the corresponding unresolved region, including the *β*20 strand, was reconstructed in a C-tail-displaced state (see **Methods** for further details). We also used these simulations to assess whether the model remained stable in a Fab30-free condition.

Both systems displayed a limited conformational plasticity throughout the simulations and showed distinct interaction signatures consistent with their respective basal and V2Rpp-bound states. Notably, basal arrestin-2 was able to preserve its autoinhibited architecture, i.e., the interactions of the “canonical polar core” (D26, R169, D290, D297 and R393) and “three-element network” (e.g., K10–D390, R7–E389) remained persistent across the simulation time [30, 31], with the latter one representing a pre-condition to maintain a close association between the C-terminal tail and the N-domain (Fig. 2A). Additional contacts between *β*1 and *β*20 further stabilised the C-tail-bound arrangement.

**Figure 2:**
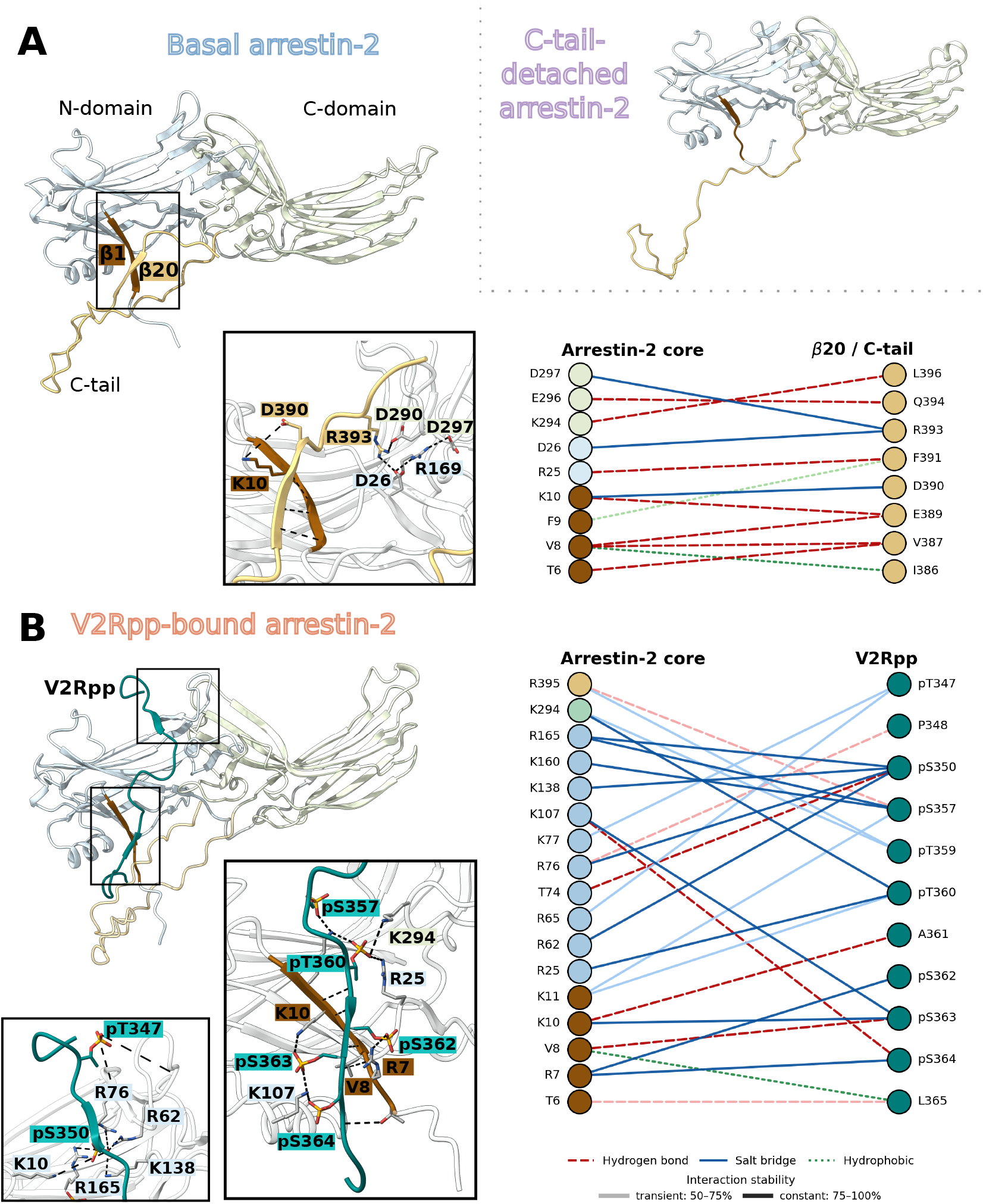
Structural representation of arrestin-2 systems investigated in this study. **(A)** Apo arrestin-2 in its basal, autoinhibited conformation. The enlarged view highlights the interaction network stabilizing the polar core and the *β*1 - *β*20 interface responsible for anchoring the C-terminal tail to the N-domain. The C-tail-detached arrestin-2 model, representing a peptide-free structure in which *β*20 is detached from the N-domain, is also shown. **(B)** V2Rpp-bound arrestin-2, showing V2Rpp engaged with the N-domain. The enlarged view highlights the principal protein–peptide interactions, stabilizing the bound state. For the basal and V2Rpp-bound systems, the corresponding interaction diagrams summarise persistent contacts identified across the simulations. Structural elements are coloured consistently across the three systems: the N-domain is shown in pale blue, the C-domain in pale green, the C-tail including *β*20 in pale gold, *β*1 in brown, and V2Rpp in dark teal.

The V2Rpp-bound simulations exhibited a different interaction network (Fig. 2B). Contacts involving C-tail residue R393 were absent or reduced, as were polar-core contacts involving D290 and D297. Only the D26–R169 contact remained persistent, showing that this part of the polar core contact network is preserved even after phosphopeptide binding. The binding of V2Rpp was stabilised by interactions between phosphorylated residues and positively charged amino acids in arrestin-2’s N-domain, such as R7, K10, K11, R62 and R76. V2Rpp binding disrupted the finger-loop ionic-lock network formed by R62–D67 and R76–D69. Indeed, these contacts fluctuated in the basal ensemble but were further weakened upon phosphopeptide binding, accompanied by more heterogeneous finger-loop orientations consistent with activation-associated reorganization (see Supplementary Fig. S1).

### 2.2 V2Rpp binding broadens the accessible rotation range beyond C-tail detachment

Next, we used the conventional MD trajectories to assess how the interdomain orientation of arrestin-2 evolved in the basal and V2Rpp-bound models. We started by measuring rotation of the C-domain relative to the N-domain following the definition used by Latorraca et al. [20] with the basal structure as the 0° reference. The V2Rpp-bound reference structure exhibited an interdomain rotation of approximately 21° relative to this structure (Fig. 3A, see **Methods** for further details). All four basal trajectories remained predominantly near 0°, with occasional fluctuations towards 10°. Conversely, the four V2Rpp-bound trajectories departed from the starting orientation (*∼*21°) and rarely returned to this initial range. Among them, during the final 5 *µ*s, two simulations sampled rotational values centred near 0°, while the other two remained centred at an average value of *∼*10° (Fig. 3B). Nevertheless, the limited number of transitions across the sampled angular range meant that equilibrium populations could not be reliably estimated using conventional MD alone.

**Figure 3:**
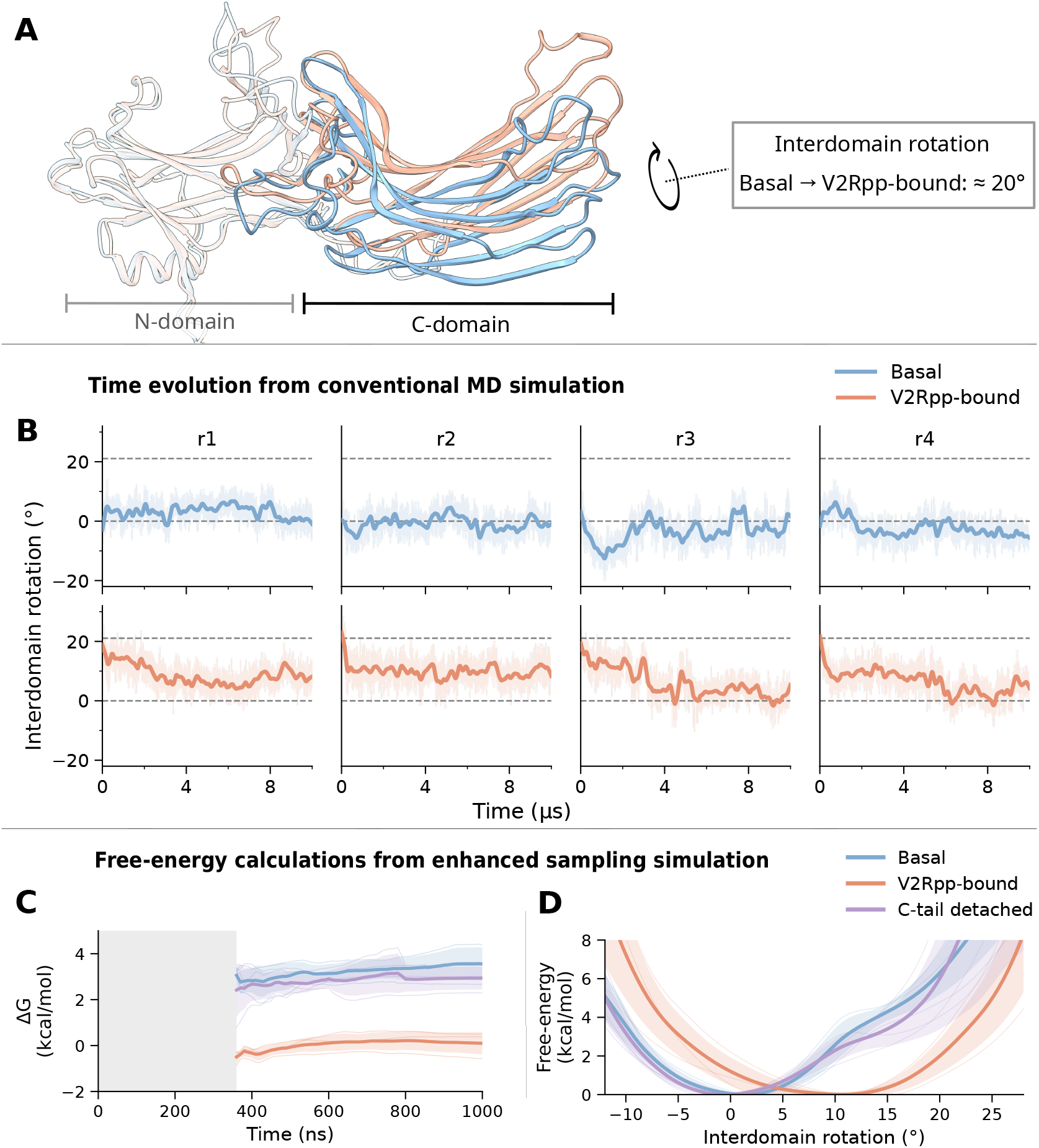
Interdomain rotation in arrestin-2. **(A)** Structural superposition of basal (blue) and V2Rpp-bound (orange) arrestin-2, aligned on the N-domain, illustrating the approximately 21*°* interdomain rotation associated with receptor engagement. **(B)** Interdomain rotation during four independent 10-*µ*s conventional MD simulations (r1–r4) of basal (top) and V2Rpp-bound (bottom) arrestin-2. Thin transparent traces show the rotation sampled at 1-ns intervals, while thick lines show smoothed trends. Grey dashed lines indicate the starting rotations of the basal (0°) and V2Rpp-bound (21°) reference structures. **(C)** Convergence of the free-energy difference between low- and high-rotation basins obtained from enhanced sampling simulations. The shaded region indicates the initial equilibration period (350 ns) excluded from the analysis. **(D)** Reweighted free-energy profiles along the interdomain rotation angle for basal, V2Rpp-bound and C-tail-detached arrestin-2. For **(C)** and **(D**), each independent trajectory is shown as a thin solid line, whereas their average is represented as a thicker solid line. The transparent areas indicate their standard deviation.

To determine the relative energetic accessibility of different interdomain orientations, we used enhanced-sampling simulations. We considered three systems: basal arrestin-2, V2Rpp-bound arrestin-2, and C-tail-detached arrestin-2. The third system allowed us to distinguish the effect of V2Rpp binding from that of C-tail detachment alone. Sampling was performed using an RMSD-based path collective variable (*s*_path_) defined by the basal and V2Rpp-bound reference structures [32]. For each system, we carried out three independent 1-µs simulations. After discarding the first 350 ns, we used the remaining 650 ns for thermodynamic analysis. To assess sampling quality, we monitored the time evolution of the main CV (see Supplementary Fig. S2). Because *s*_path_ describes the broader conformational transition, we subsequently projected the trajectories onto the interdomain-rotation angle to obtain more readily interpretable free-energy profiles. Among rotational states, the freeenergy differences reach stable plateaus (Fig. 3C), supporting the reliability of the estimates for each simulation. Likewise, each individual profile follows the same trend along the analysed CV space (Fig. 3D). The three independent simulations produced consistent free-energy profiles (Fig. 3C,D). Basal arrestin-2 strongly favoured orientations close to the basal reference state: its free-energy increased progressively as the interdomain rotation approached the V2Rpp-bound reference angle of 21°. In contrast, the V2Rpp-bound landscape was substantially flatter, with a shallow minimum near 10°. The *≃*21° orientation observed in the reference structure was therefore accessible but not strongly favoured. Thus, V2Rpp binding broadens the range of energetically accessible interdomain orientations rather than stabilizing a single highly rotated state.

C-tail detachment alone did not reproduce this effect. In the absence of V2Rpp, C-tail-detached arrestin-2 retained a free-energy profile similar to that of basal arrestin-2, with a minimum near 0° and increasing free-energy towards 21° (Fig. 3D). These results indicate that C-tail detachment alone is insufficient to explain the effect of V2Rpp binding on the free-energy profile of interdomain rotation. Nevertheless, compared with basal arrestin-2, C-tail detachment lowered the free-energy penalty at larger *s*_path_ values, i.e., conformations toward the V2Rpp-bound reference. The shift was limited: this region remained energetically costly to access in both basal and C-tail-detached arrestin-2, unlike in V2Rpp-bound arrestin-2. The difference persisted at comparable interdomain rotation angles (see Supplementary Fig. S3), indicating that C-tail detachment affects structural features beyond the interdomain twist.

### 2.3 C-tail detachment is unfavourable in the absence of the phosphopeptide

The basal and C-tail-detached systems also differ along conformational coordinates that are not captured by interdomain rotation alone (see Supplementary Fig. S3). This raises a separate question: how energetically accessible is spontaneous C-tail detachment in basal arrestin-2? To address this, we calculated the free-energy cost of detaching *β*20 from its basal binding site in the absence of V2Rpp. We used a dedicated OneOPES enhanced-sampling protocol in which *β*20 displacement was sampled within a cylindrical restraint (Fig. 4A).

**Figure 4:**
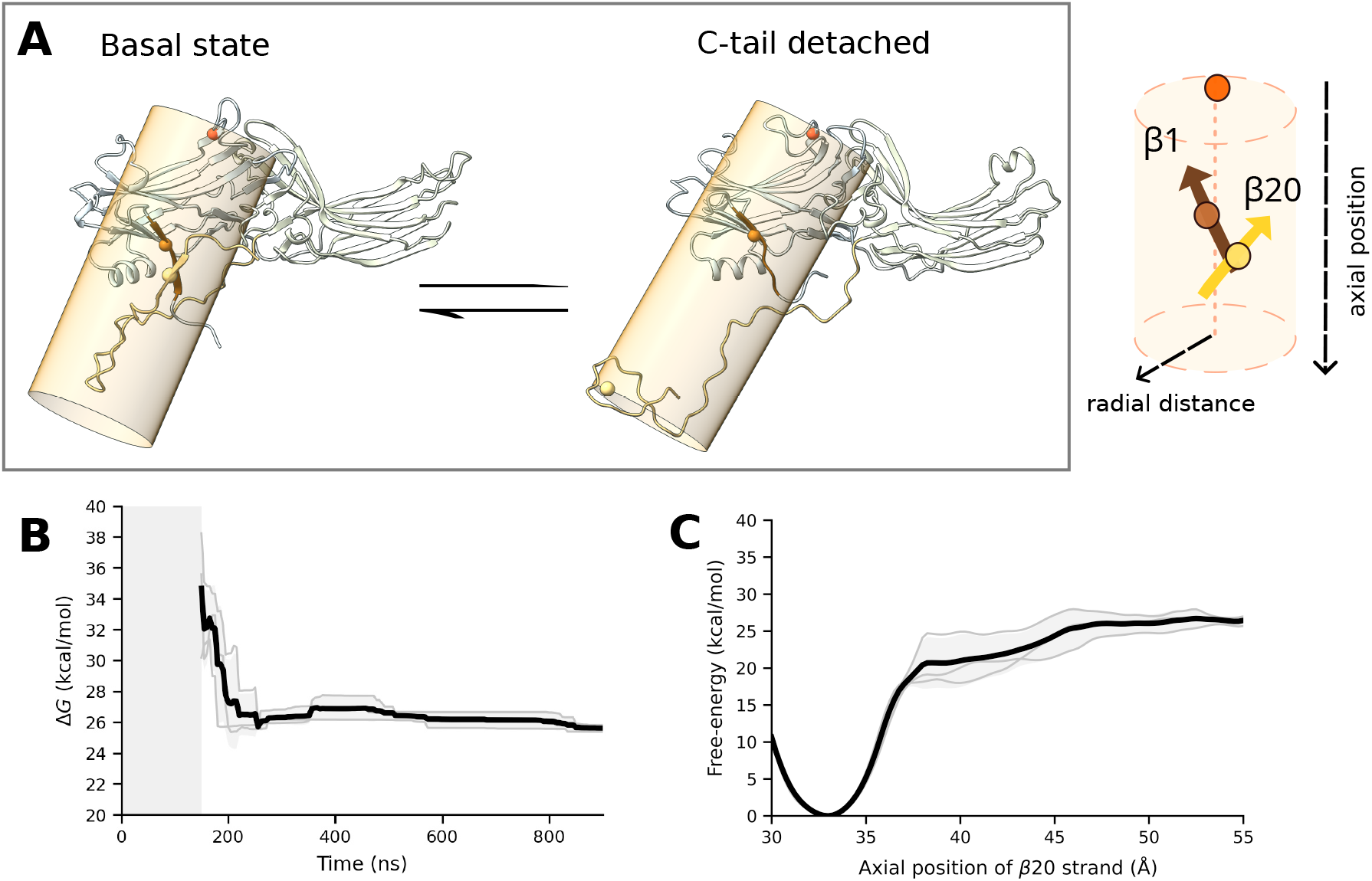
Free-energy landscape describing detachment of the arrestin-2 C-terminal tail from the autoinhibited state in the absence of the V2Rpp. **(A)** Representative structures of the basal, C-tail-attached state (left) and the C-tail-detached state (right), between which the free-energy difference was evaluated. Inset: schematic of the cylindrical-restraint geometry. The reference axis is defined by two anchor points in the N-domain: the centre of mass of the finger loop (orange) and the *β*1 strand (brown). The position of the *β*20 strand (yellow), which forms the N-domain protein–protein interaction anchor point in the autoinhibited state, is described relative to this axis by its axial position (the projection along the axis) and radial distance (the orthogonal distance from the axis). C-tail detachment is therefore monitored as the movement of *β*20 away from the *β*1 anchor point. See Methods for details. **(B)** Convergence of the free-energy difference between the attached and detached states over simulation time; the shaded region marks the initial 150 ns equilibration period excluded from the analysis. **(C)** Free-energy profile for *β*20 strand displacement along the axial position. For **(B)** and **(C)**, each independent trajectory is shown as a thin solid line, whereas their average is represented as a thicker solid line. The transparent shaded areas indicate their standard deviation.

The detachment simulations were biased using a set of CVs designed to capture complementary aspects of C-tail disengagement. The final detachment profile was analysed along an axial projection coordinate (*pp*.*proj* hereafter), which tracks *β*20 displacement along the sampled detachment direction (see **Methods** for further details). We collected three independent 900 ns trajectories, analyzing the final 750 ns after discarding the initial out-of-equilibrium portions. To assess sampling quality, we monitored the time evolution of the main CVs (see Supplementary Fig. S4) as well as the behavior of *pp*.*proj* (see Supplementary Fig. S5). These analyses showed that both C-tail-bound and C-tail-detached configurations were extensively explored.

All three replicas yielded a large free-energy cost for *β*20 detachment (Δ*G*_*unbinding*_ of 25.6 *pm* 0.2 )kcal/mol (Fig. 4B). Because the calculation follows a defined, restrained detachment pathway, this value should be interpreted as the effective energetic cost of *β*20 displacement along that pathway rather than as an absolute binding free-energy. To test whether the cylindrical restraint artificially increased this cost, we repeated the analysis while allowing the detached C-tail full positional freedom. The free-energy difference decreased by only 1 kcal/mol, indicating that the large energetic penalty is not primarily caused by the restraint. These simulations therefore consistently indicate that *β*20 detachment is strongly unfavourable in basal arrestin-2 under the sampled conditions.

To better discern the structural details associated with the unbinding, we re-projected the free-energy as a function of both *pp*.*proj* and multiple meaningful intramolecular interactions within arrestin-2 (i.e., D26–R393, D290–R393, K10–R393, R7–D390, and D26–R169). As expected, in all five free-energy surfaces, the deepest minimum consistently coincided with the *pp*.*proj* value corresponding to the attached configuration of *β*20 strand (*pp*.*proj ∼*30–35 Å). In this region, the basal C-tail-associated interaction network adopted its most compact configuration, including contacts directly linking the C-terminal tail to the N-domain, such as D26–R393, K10–R393, and R7–D390, as well as the lariat-loop-associated D290–R393 contact (see Supplementary Fig. S6A-D).

In particular, Fig. S6A and Fig. S6B show a two-step unbinding mechanism, where both D26–R393 and D290–R393 distance must break before the full *β*20 detachment takes place. Conversely, the K10–R393 and R7–D390 contacts weakened more gradually along the sampled pathway (see Fig. S6C-D). Lastly, the salt bridge D26–R169 remained comparatively stable over a broad range of projection-coordinate values, suggesting that this component of the polar core is largely preserved during C-terminal tail detachment (see Supplementary Fig. S6E). Moreover, the persistence of D26–R169 indicates that the detachment pathway preserves the internal organization of the arrestin core. This supports the interpretation that the observed free-energy changes arise primarily from the disruption of the C-tail-associated interaction network rather than from global destabilization of the protein. This behaviour is also consistent with unbiased MD simulations, in which the D26–R169 interaction remained stable despite fluctuations in the C-terminal tail.

## 3 Discussion and conclusions

Arrestin activation involves several coupled structural changes, including binding of the phospho-rylated receptor tail, displacement of the arrestin C-tail, rearrangement of basal intramolecular interaction networks and changes in the relative orientation of the N- and C-domains [33, 34, 35, 36]. In this study, we investigated whether interdomain rotation in tail-engaged arrestin-2 is due to the phosphopeptide binding itself or rather arises from the detachment of the autoinhibitory C-tail. To address this question, we measured the interdomain-rotation landscape (and how it is shaped by V2Rpp binding), complementing our investigation with the estimate of the energetic cost associated to C-tail detachment in the absence of a bound phosphopeptide.

In basal arrestin-2, the free-energy profile showed a preference for interdomain-rotation angles in the close proximity to the basal reference structure, whereas larger rotations towards the V2Rpp-bound reference are energetically discouraged. C-tail-detached arrestin-2 demonstrated a similar preference for *∼*0° interdomain-rotation angles. This indicates that C-tail detachment alone is insufficient to account for the interdomain-rotation landscape observed upon phosphopeptide binding. This behaviour is consistent with earlier simulations in which C-tail removal permitted larger rotations but did not make strongly rotated conformations dominant [20].

V2Rpp binding had a qualitatively different effect. Even in the absence of receptor-core interactions and Fab30, V2Rpp broadened and flattened the interdomain-rotation free-energy landscape. Rather than stabilizing a single rotated conformation, V2Rpp reduced the energetic differences among several basal-like and rotated states. The 21° orientation seen in the V2Rpp-bound crystal structure remained accessible but was not a dominant free-energy minimum. We therefore interpret this structure as one member of a broader V2Rpp-bound conformational ensemble rather than as the uniquely preferred state.

This interpretation is also consistent with experimental evidence that Fab30 shifts arrestin’s conformational equilibrium and stabilises the V2Rpp complex, supporting a model in which the antibody enriches a more strongly rotated conformation within a broader V2Rpp-bound ensemble [28, 37]. Moving on, the calculation of the unbinding free-energy provides a perspective on C-tail release in the absence of phosphopeptide binding. The high free-energy cost of *β*20 unbinding suggests that C-tail-detached conformations are only scarcely populated by basal arrestin-2 under the sampled conditions. This is qualitatively consistent with smFRET evidence for strong basal C-tail autoinhibition [37], although it contrasts with a basal-state NMR exchange process interpreted as C-tail detachment [38]. Whether the NMR-inferred state and the detached *β*20 configurations sampled here correspond to the same form of C-tail disengagement remains unresolved. Our calculation therefore reports on the free-energy difference between the attached state and the displaced *β*20 state defined within the sampled detachment framework. It should be interpreted as the energetic cost of removing the C-tail from its basal interface along this defined unbinding coordinate, rather than as a description of all possible modes of C-tail dynamics or exchange.

Tail-engaged receptor–arrestin complexes can support particular endocytic and signaling responses without core engagement, although these observations do not establish which interdomain orientations are populated [27, 21]. The range of interdomain orientations accessible in our soluble V2Rpp model leaves scope for receptor-core, membrane or effector interactions to favor particular orientations. Whether such population shifts contribute to the functional properties of tail-engaged complexes remains to be established.

A remaining question is whether the broadening of the interdomain-rotation landscape observed here is specific to V2Rpp or represents a general consequence of the engagement of other receptor phosphotails. This work provides a framework for future studies investigating how different phosphorylation patterns influence arrestin recruitment.

## 4 Methods

### 4.1 Systems preparation

The initial coordinates for basal arrestin-2 were derived from the chain A of the crystal structure PDB ID: 8AS4 [39]. The missing regions (residues 1-4; 68-71, 92-94, 134-137 and 358-385) were reconstructed by incorporating the corresponding segments from the AlphaFold2 (AF2) model after structural alignment performed with ChimeraX [40, 41]. In available crystal structures of basal arrestin the finger loop usually adopts bent conformation oriented toward the N-domain and participates in maintaining the ionic lock that stabilise the inactive state. Reconstruction of the finger loop using the AF2 model resulted in a more upright initial orientation of the loop, resembling a more open arrangement. Such an arrangement was treated solely as a starting geometry. As no positional restraints were applied to this region during the production runs of the unbiased MD simulations, the loop was free to sample conformational space, including conformations resembling the bent state.

In particular, the *proximal* C-terminal region (residues 358-385) was retained to preserve structural continuity around the C-tail autoinhibitory element, i.e, the region containing the *β*20 strand.

A similar strategy was employed to model the tail-engaged arrestin-2 in complex with the V2Rpp. In detail, the model for tail-engaged arrestin-2 was extracted from the crystal structure PDB ID: 8AS3 [39], which contains the arrestin-2 in complex with the 6P CCR5 phosphopeptide (CCR5pp). The unresolved regions of 8AS3 (residues 1–5; 65–69; 309–312; and 358–onward) were reconstructed by incorporating the missing segments from the arrestin-2/CCR5pp complex generated using AF2 [42]. To be consistent with the basal model, we did not include the *distal* C-terminal tail, trimming the primary sequence of our tail-engaged arrestin-2 to amino acid 396. V2Rpp was introduced by superimposing the V2Rpp-bound arrestin structure (PDB ID: 4JQI [18]) onto the reconstructed 8AS3 model and retaining only the coordinates of the bound phosphopeptide. Missing peptide fragments (i.e., Q354 and D355) were modeled using the “Model Loops” tool in ChimeraX. The basal and tail-engaged systems were then prepared for subsequent MD simulations using the CHARMM-GUI solution builder [43]. Both V2Rpp and arrestin-2 were parametrised with the CHARMM36m force field [44, 45], with C-terminus of the latter protected by a methylamide. The proteins were embedded within a rectangular periodic simulation box and solvated with explicit TIP3P water molecules (pH 7.0; salinity 0.15 M NaCl).

### 4.2 Unbiased MD simulations

To equilibrate the basal state of arrestin-2, a stepwise heating protocol with progressively reduced positional restraints on the protein’s heavy atoms was applied to relax potential structural artifacts. Equilibration consisted of consecutive 500 ps NPT and 500 ps NVT simulations at increasing temperatures (100-300 K) in 50 K increments. After the NVT simulation at 300 K, the final NPT equilibration was run for 5 ns. For the tail-engaged structure, equilibration was performed using the same thermal ramp but with a two-stage restraint detachment scheme to stabilise the initial phosphopeptide binding. During the first stage, the phosphopeptide was kept under strong positional restraints, whereas the protein restraints were gradually reduced. In a subsequent stage, positional restraints on the phos-phopeptide were progressively released, and the final NPT equilibration was run for 5 ns. Covalent bonds involving hydrogen atoms were constrained using LINCS [46, 47], allowing an integration time step of 2 fs. Short-range Coulomb and van der Waals interactions were truncated at 1.0 nm, and long-range electrostatics were treated with the particle-mesh Ewald (PME) method [48]. The temperature was maintained at 300 K using the velocity-rescale (V-rescale) thermostat [49], and the pressure was controlled at 1 bar using the isotropic C-rescale barostat [50].

Following equilibration, all positional restraints were removed and unbiased MD simulations were performed. For both states, four independent 10-*µ*s trajectories were generated, yielding a total of 40 *µ*s of simulation per system (80 *µ*s in total). Replicas were initiated from the equilibrated structures using initial velocity distributions sampled from the Maxwell-Boltzmann distribution at 300 K. Trajectories were run in the NPT ensemble at 300 K and 1 bar under periodic boundary conditions. The same force field parameters, thermostat, barostat, and non-bonded interaction settings as described above were used during the production simulations. Both equilibration and production runs have been carried out through GROMACS 2023 [51].

### 4.3 Analysis of trajectories and molecular contacts

All trajectories were re-imaged to remove periodic boundary condition artifacts, and the systems were centred in the simulation box for further analysis with GROMACS tools. Unless otherwise specified, the trajectories were structurally aligned using C*α* atoms of *β*-strands forming the N-domain, which served as a stable structural reference frame. This alignment strategy has also been applied in previous works [20, 33, 52, 53].

Interaction analyses were performed using in-house Python scripts based on the MDAnalysis library [54, 55]. For each frame, molecular contacts were identified. Electrostatic interactions were analysed, with particular focus on salt bridges. A salt bridge was assigned when the distance between the charged groups was shorter than 4.5 Å. Hydrogen bonds were identified using a geometric definition that required a donor–acceptor distance shorter than 3.5 Å and a donor–hydrogen–acceptor angle greater than 120°. The donor–hydrogen covalent distance was also constrained to 1.2 Å to ensure proper donor assignment.

### 4.4 Analysis of finger loop dynamics

The orientation of the finger loop relative to arrestin was quantified using geometry-based CVs calculated with PLUMED 2.9 [56, 57]. The principal axis of the protein was defined as the vector that connects the centers of mass (COMs) of arrestin’s N- and C-domains. The domain COMs were calculated using C*α* atoms belonging to residues forming *β*-strands in each domain. The orientation of the finger loop was described by a vector that connects the COMs of the residues involved in the “ionic lock” of the finger loop. The ionic lock corresponds to a set of contacts between evolutionarily conserved negatively charged residues (D67 and D69) and two conserved arginines (R62 and R76) located in the *β*-strands S5 and S6 of the N-domain.

The angle between the principal axis of the protein and the finger loop was calculated using a four-point definition corresponding to the angle between the two vectors. Angles close to 90° indicate an orientation of the finger loop approximately perpendicular to the principal axis of the protein, while deviations from this value indicate changes in the positioning of the loop relative to this axis. To evaluate the directional preference of loop orientation, a signed projection of the finger loop vector onto the principal axis was additionally computed. Values with the negative sign indicate the direction towards the N-domain, while positive values indicate the direction towards the C-domain.

### 4.5 Enhanced sampling of interdomain rearrangements

Interdomain rearrangements were sampled using a two-reference, RMSD-based path collective variable (path-CV, *s*_path_) connecting the basal and V2Rpp-bound conformations. The metric was calculated from C*α* atoms in structured elements shared by the two references [32, 58, 59, 60]. N-domain atoms were used for structural alignment, thereby removing overall translation and rotation, whereas the displacement metric included structured atoms from both domains. The progress variable *s* therefore reports movement along the broader structural transition between the references: values close to 0 are basal-like, whereas values close to 1 approach the V2Rpp-bound reference. Because this coordinate also includes local and collective rearrangements that accompany domain rotation, it was not interpreted as a rotation angle. A restraint on the orthogonal path variable *z* limited sampling of conformations distant from both reference nodes.

Sampling was enhanced using a OneOPES-like multiple-replica scheme [61]. The protocol differed from standard OneOPES because the OPES Expanded multithermal thermal ramp was not necessary for achieving the targeted sampling quality. For each of the basal, C-tail-detached, and V2Rpp-bound systems, three independent calculations were performed, each comprising one convergence-dedicated replica and three progressively more exploratory replicas. All four replicas were biased along *s*_path_; replicas 1–3 also used auxiliary biases. The complete biasing scheme for all replicas is provided in the **Supplementary Table S1**. The free-energy analyses reported here used the convergence-dedicated replica from each independent calculation. Each trajectory was 1 *µ*s long. The first out-of-equilibrium 350 ns were discarded, and the remaining 650 ns were reweighted at 300 K using both the OPES path bias and the *z*-wall bias recorded for each frame.

### 4.6 Projection onto interdomain rotation and free-energy analysis

To isolate the rotation component of the sampled rearrangement, trajectories were projected onto a signed planar interdomain-rotation angle adapted from the analysis introduced by Latorraca and coworkers [20]. For each frame, independent rigid-body fits of the N- and C-domain C*α* selections were used to determine the relative orientation of the two domains. The resulting rotation was projected onto the inactive-to-active rotation axis defined by the basal and V2Rpp-bound reference structures used in this study and was reported in degrees. The basal reference is therefore close to 0°, whereas the V2Rpp-bound reference is approximately 20°. No loop, C-tail, peptide or solvent atoms entered this geometric measurement. One- and two-dimensional free-energy surfaces were calculated separately for the three independent simulations of each system and then summarised across replicates. The one-dimensional profiles report the replicate mean, with the between-replicate standard deviation shown as a shaded band. A joint projection against the *s*_path_ was used to assess how the focused rotation angle relates to broader inactive-to-active structural progress. The projected rotational profiles describe the conformations sampled within the restrained path region.

### 4.7 Building bio-inspired collective variables for the structural rearrangement associated with the detachment of the autoinhibited state

To identify collective variables describing arrestin’s C-tail detachment, two structural reference ensembles corresponding to the end states of the conformational transition were required. Following strategies used in the recent study of protein-folding [62], where discriminative features are identified by comparing distinct structural states, two ensembles representing the autoinhibited and detached states of arrestin-2 were generated. In the basal state, *β*20 strand of the C-terminal tail remains positioned against *β*1 of the N-domain, forming an intramolecular interface that stabilises the inactive state. Therefore, separation of *β*20 from *β*1 was treated as the primary conformational change.

The autoinhibited ensemble was obtained from an unbiased MD simulation of the basal state. During this simulation, additional geometrical confinement was applied to prevent the detachment of the flexible C-terminal tail into the bulk solvent. The confinement was implemented in PLUMED 2.9 using a cylindrical restraint based on a projection-on-axis coordinate. The axis was constructed using two reference pseudoatoms located within the *β*-sheet core of the N-domain. The position of *β*20 strand was represented by the geometric center of C*α* atoms from residues 384-392. Its position relative to the axis was decomposed into a projection along the axis (*pp*.*proj*) and an orthogonal distance from the axis (*pp*.*ext*). A soft upper-wall potential was applied to the orthogonal component, restricting the radial detachment of the C-terminal segment within a cylinder of 1.5 nm radius, without constraining motion along the axis.

To obtain the detached-state ensemble, *β*20 was displaced away from *β*1 using steered MD (SMD). The steering procedure served only to overcome the autoinhibitory contacts and generate a geometry with the detached C-terminal tail. The structure obtained during SMD simulation was subsequently subjected to unbiased MD simulation. During the simulation, additional geometrical confinement based on the same projection-onto-axis coordinate was applied to prevent spontaneous reattachment of *β*20 to the *β*1. Soft wall potentials were applied to both components of the coordinate: the radial upper wall was maintained as in the autoinhibited state, whereas a lower wall was introduced along the axial projection. The lower wall was positioned at 6.5 nm from the axis origin. Additionally, a soft RMSD restraint was used on the C*α* atoms of the *β*-strands 1-19 to preserve the overall fold of arrestin-2.

The resulting trajectories served as input for automatic feature identification using the *bioin-spired_features*.*py* script (available at: https://github.com/valeriorizzi/FoldingFeatures) [63]. This procedure yielded two complementary CVs. The first CV captures high-energy microscopic interactions by monitoring hydrogen-bond patterns that distinguish intramolecular protein–protein hydrogen bonds from protein–water interactions and also incorporates angular information. The second CV describes mesoscale organization by quantifying side-chain contacts, thereby reporting on the compactness of the interface between the analysed *β*1 and *β*20 strands.

### 4.8 Biased MD simulations of the transition from the autoinhibited state

The OneOPES strategy [61] was employed to enhance sampling of the detachment of the flexible C-terminal tail, specifically the detachment of *β*20 of the C-terminal tail from *β*1 of the N-domain. In each of the three independent simulations, eight replicas were executed: one convergence-dedicated replica (**replica 0**) and seven exploratory replicas (**replicas 1-7**). All replicas were biased by OPES Explore [64] as the main sampling bias, while the exploratory replicas were progressively heated to 335 K with OPES Expanded [65] to help overcome hidden degrees of freedom.

The main bias acted on three CVs and was constructed using two bio-inspired features described above and an additional structural separation CV. Specifically, the biased variables were: (i) a hydrogen-bond–based CV describing formation and disruption of inter-strand hydrogen-bond networks, (ii) a side-chain packing CV reporting on the compactness of the *β*1/*β*20 interface, and (iii) a structural separation CV defined as a distance between the geometric center of *β*20 and the geometric center of C*α* atoms forming *β*-strands 1–19 of protein [62]. This primary bias was present in all replicas. Water-coordination CVs were also applied in all replicas to enhance the sampling of local hydration and solvent exposure at the *β*1 interface [66, 67, 68]. Further water-coordination CVs were introduced progressively across replicas to improve the sampling of selected local interactions during the *β*20 displacement. The complete biasing scheme for all replicas is provided in the **Supplementary Table S3**. In all replicas, a set of restraints was introduced to confine sampling to physically relevant regions of the conformational space. An RMSD wall was applied to the C*α* atoms of the *β*-sheet core to preserve the global fold of arrestin-2 while allowing local rearrangements. A restraint on the orthogonal extension component, *pp*.*ext*, limited excessive lateral displacement of the *β*20 strand. In parallel, a wall applied along the axial projection coordinate, *pp*.*proj*, prevented drift across periodic boundaries along the detachment direction [69, 70, 71]. An additional restraint on the structural separation coordinate maintained separation from the autoinhibited interface once detachment occurred. Finally, restraints coupling the coordinate of the axial projection with the hydrogen-bond and side-chain packing variables ensured that the separation of *β*20 strand corresponded to an actual loss of interstrand contacts. The resulting trajectories were reweighed and projected onto *pp*.*proj*, monitoring *β*20 displacement along the sampled detachment pathway.

## Acknowledgements

This work was financially supported by the Swiss National Science Foundation funding schemes (project numbers: 200021_204795 and CR00-5-239843). We acknowledge the Swiss National Super-computing Centre for supercomputer time allocations on Piz Daint and Alps (project IDs: s1274, lp84, and lp166). We thank Valerio Rizzi, Nicola Piasentin, and Anton Hanke for valuable discussions on the construction and optimization of the collective variables. We also thank Stephan Grzesiek for insightful discussions, which offered a valuable perspective based on his experience with NMR studies of arrestin. We thank Naomi Latorraca for sharing her original script for monitoring the interdomain rotation in arrestins and for useful advice.

