## Supplementary Material for "Vasopressin V2 receptor phosphopeptide reshapes the conformational landscape of arrestin-2 beyond C-tail detachment"

### Supplementary Data 1

To characterize the conformational response of the finger-loop region in basal and V2Rpp-bound arrestin-2, we monitored a set of dedicated structural descriptors calculated from the unbiased MD trajectories: two ionic lock distances, a finger-loop angle and a domain-projection coordinate. The ionic-lock distances monitored the R62—D67 and R76—D69 residue pairs, reporting on local rearrangements of the finger-loop anchoring region. The arrestin principal axis was defined between the N- and C-domain centers of mass, whereas the finger-loop vector was defined using residues of the conserved ionic lock region. The angle and signed projection of this vector relative to the N–C domain axis report on finger loop orientation and directional preference toward the N- or C-domain, respectively.

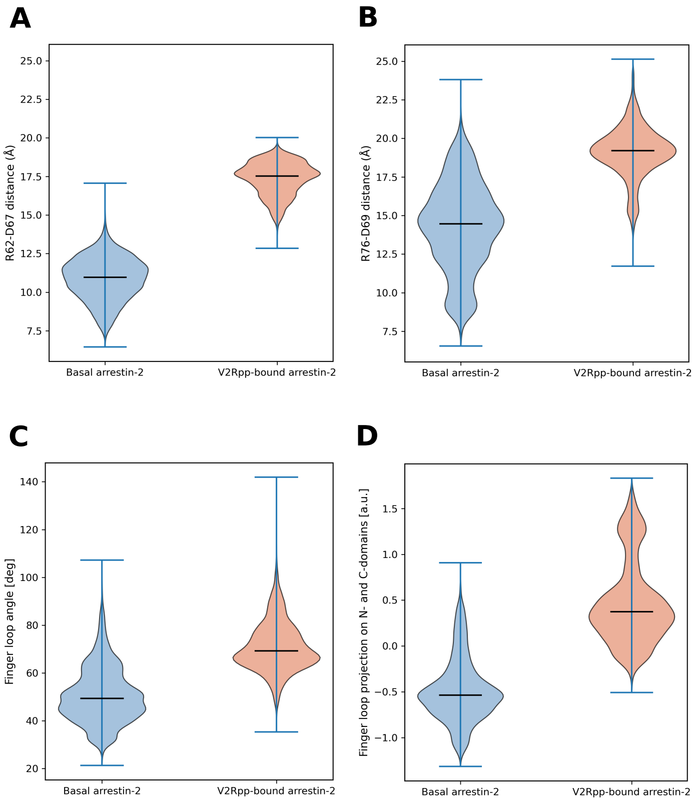


**Figure S1: Distributions of finger loop descriptors calculated from unbiased molecular dynamics (MD) simulations of basal and V2Rpp-bound arrestin-2. (A, B)** Distances between the ionic-lock residue pairs R62–D67 and R76–D69. **(C)** Distribution of the finger-loop angle. Angles close to (90°) indicate an orientation of the finger loop approximately perpendicular to the principal axis of the protein, while deviations from this value indicate changes in the positioning of the loop relative to this axis. **(D)** Projection of the finger loop orientation onto the N- and C-domain reference directions. Negative values indicate orientation towards the N-domain, whereas positive values indicate orientation towards the C-domain. Violin plots show the sampled distributions with horizontal lines indicating median values. Compared with basal arrestin-2, the V2Rpp-bound state samples larger ionic lock distances and higher finger loop angles, consistent with reorganization of the finger loop region upon phosphopeptide binding.

### Supplementary Data 2

To investigate the interdomain rotation landscape of arrestin-2, we used an enhanced sampling setup based on a main OPES Explore bias applied to an RMSD-based path collective variable, $s_{\mathrm{path}}$, connecting the basal and V2Rpp-bound conformations. In replicas 1-3, additional biases were applied to improve sampling of structural features coupled to arrestin activation, including including rearrangements of the polar core and finger-loop ionic lock, interdomain/finger-loop orientation, and contact-map changes. The OPES Explore barrier for the main CV was set to 50 kJ/mol whereas the barriers for the interdomain/finger-loop orientation, polar-core rearrangement and contact-map CVs were set to 10, 5, and 5 kJ/mol, respectively.

**Table S1:** Biasing scheme used for the interdomain rotation simulations. All replicas (0-3) were biased along the RMSD-based path collective variable $s_{\mathrm{path}}$. Replicas 1–3 additionally included polar-core distances R169–D290 and R169–D26, finger-loop ionic-lock distances R62–D67 and R76–D69 and the finger-loop orientation angle $\theta_{\mathrm{FL}}$ between the principal axis of arrestin-2 and the finger loop. Replica 3 further included apo- and holo-specific contact-map variables.

| Replica | Main CV | Additional biased CVs |
|---|---|---|
| 0 | $s_{\mathrm{path}}$ | – |
| 1 | $s_{\mathrm{path}}$ | $\mathbf{d}_{\mathrm{FL-lock}},\ \theta_{\mathrm{FL}},\ \mathbf{d}_{\mathrm{PC}}$ |
| 2 | $s_{\mathrm{path}}$ | $\mathbf{d}_{\mathrm{FL-lock}},\ \theta_{\mathrm{FL}},\ \mathbf{d}_{\mathrm{PC}}$ |
| 3 | $s_{\mathrm{path}}$ | $\mathbf{d}_{\mathrm{FL-lock}},\ \theta_{\mathrm{FL}},\ \mathbf{d}_{\mathrm{PC}},\ \mathbf{c}_{\mathrm{state}}$ |

To assess whether the main twisting CV was efficiently sampled, we monitored its time evolution in the twisting OneOPES simulations. The analyzed CV corresponds to the RMSD-based path collective variable $s_{\mathrm{path}}$, describing the conformational change along the twisting pathway. Its broad sampling over time indicates that the simulations explored different conformations along the twisting pathway across all three simulated systems.

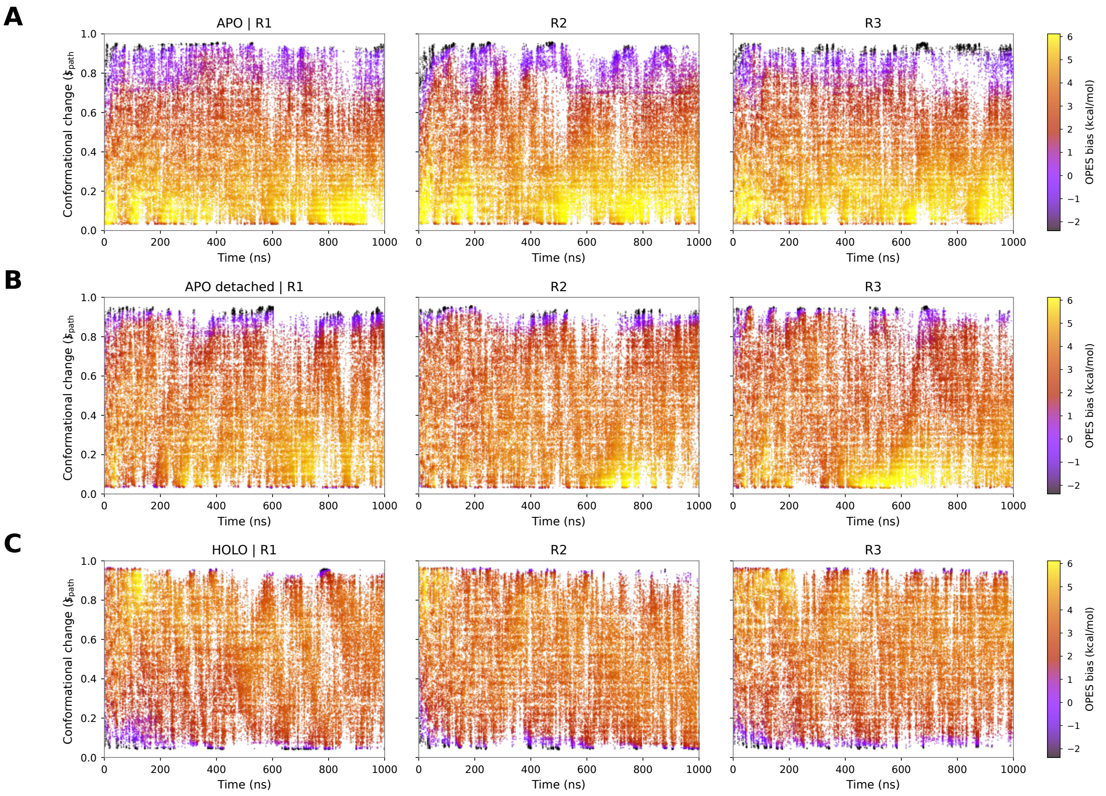


**Figure S2: Sampling of the main CV in the twisting OneOPES simulations.** Sampling of the RMSD-based path collective variable $s_{\mathrm{path}}$ describing the conformational change along the twisting pathway as a function of time for **(A)** APO, **(B)** APO detached, and **(C)** HOLO systems. For each panel, data points are shown for all three replicas (R1, R2, and R3) and colored according to the accumulated bias potential.

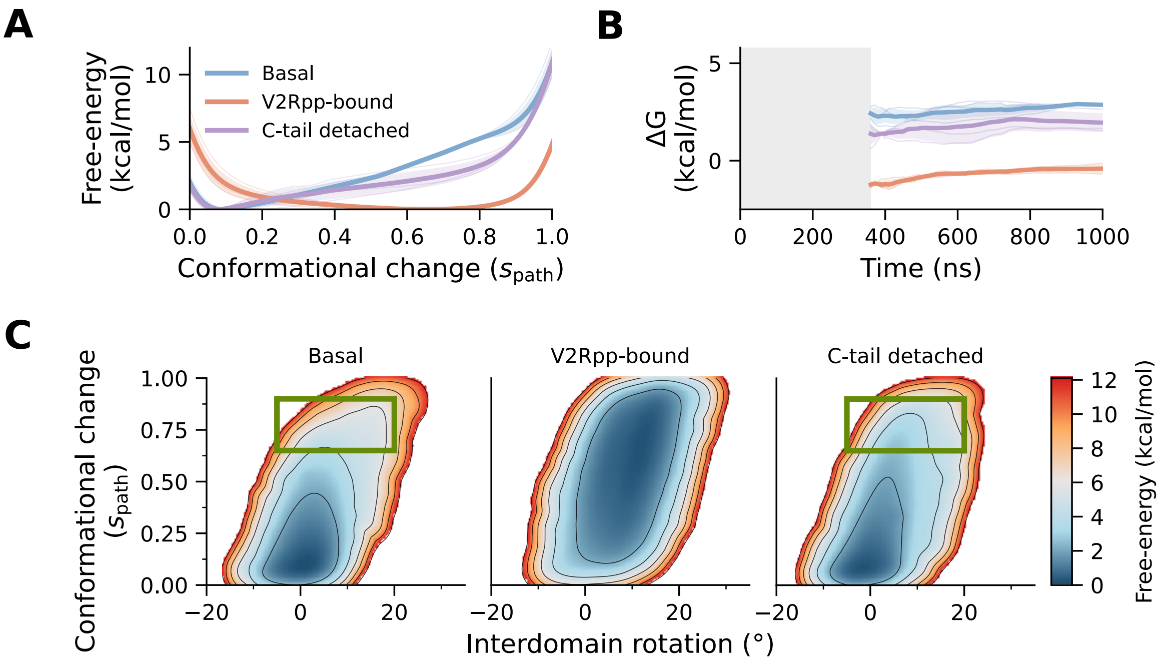


**Figure S3: Free-energy landscapes (FES) along the path collective variable (CV) used for enhanced sampling.** The RMSD-based path collective variable $s_{\mathrm{path}}$ describes conformational change between the basal and V2Rpp-bound reference structures and was used to bias the OneOPES simulations. **(A)** One-dimensional free-energy profiles along $s_{\mathrm{path}}$ for basal, V2Rpp-bound and C-tail-detached arrestin-2. **(B)** Convergence of the free-energy difference between the two regions of the path, $\Delta G$ . The grey region precedes the first estimate at 360 ns. **(C)** Two-dimensional free-energy landscapes projected onto $s_{\mathrm{path}}$ and the interdomain rotation angle. Green rectangles highlight a region in which C-tail-detached arrestin-2 has a lower relative free-energy penalty than basal arrestin-2 at comparable interdomain rotation angles. Contours are separated by 2 kcal/mol. In panels **(A, B)**, thin lines show individual calculations, thick lines their mean, and shaded regions one standard deviation.

### Supplementary Data 3

To place the simulated interdomain rotations in the context of experimentally determined structures, we compared the interdomain rotation angles of human arrestin-2 structures available in the Protein Data Bank (PDB) (Table S2)

**Table S2: Interdomain rotation angles of experimentally determined human arrestin-2 structures.**

| PDB | Chain | Experimental system | Engagement | Fab30 family | Rotation (°) |
|---|---|---|---|---|---|
| 8AS4 | B | Isolated arrestin-2 | Basal | No | 0.74 |
| 6UP7 | B | arrestin-2–NTSR1 | Core-engaged | No | 13.10 |
| 9KUW | A | arrestin-2–APLNR | Core-engaged | Yes | 15.62 |
| 9MBA | B | arrestin-2–mGlu8 | Core-engaged | Yes | 16.71 |
| 9KUX | A | arrestin-2–APLNR | Core-engaged | Yes | 17.16 |
| 9II3 | A | arrestin-2–mGlu3 | Core-engaged | Yes | 17.28 |
| 9UYH | A | arrestin-2–GPR1 (chemerin) | Core-engaged | Yes | 17.44 |
| 9KUV | A | arrestin-2–APLNR | Core-engaged | Yes | 17.53 |
| 9LXP | B | arrestin-2–PTH1R (LA-PTH) | Core-engaged | Yes | 17.54 |
| 9L8L | C | arrestin-2–$\beta_2$AR–Gs (atazanavir) | Tail-engaged | No | 17.59 |
| 9UYL | A | arrestin-2–GPR1 (chemerin) | Core-engaged | Yes | 17.71 |
| 27EM | B | arrestin-2–PAC1R (PACAP27) | Core-engaged | Yes | 18.02 |
| 8JRV | A | arrestin-2–GCGR (glucagon) | Tail-engaged | Yes | 18.03 |
| 9UYJ | A | arrestin-2–GPR1 (chemerin) | Core-engaged | Yes | 18.15 |
| 25GN | C | arrestin-2–FZD6 | Core-engaged | Yes | 18.16 |
| 8WRZ | A | arrestin-2–CB1 | Core-engaged | Yes | 18.22 |
| 6PWC | A | arrestin-2–NTSR1 | Core-engaged | Yes | 18.24 |
| 9II2 | A | arrestin-2–mGlu3 | Core-engaged | Yes | 18.36 |
| 9UYN | A | arrestin-2–GPR1 (ligand-free) | Core-engaged | Yes | 18.37 |
| 9UYI | A | arrestin-2–GPR1 (chemerin) | Core-engaged | Yes | 18.43 |
| 9LXR | B | arrestin-2–PTH1R (LA-PTH) | Core-engaged | Yes | 18.44 |
| 9LY2 | B | arrestin-2–PTH1R–V2R-tail (LA-PTH) | Core-engaged | Yes | 18.50 |
| 9M0D | B | arrestin-2–NTSR1 | Core-engaged | Yes | 18.54 |
| 8ZYT | B | arrestin-2–NTSR1 (SBI-553) | Core-engaged | Yes | 18.58 |
| 7R0J | C | arrestin-2 + V2R phosphopeptide | Phosphopeptide | Yes | 18.65 |
| 22JZ | C | arrestin-2–mGlu7 | Core-engaged | Yes | 18.77 |
| 9LY3 | B | arrestin-2–PTH1R–V2R-tail (LA-PTH) | Core-engaged | Yes | 18.91 |
| 9IJR | C | arrestin-2–GPR52 (c17) | Core-engaged | Yes | 19.14 |
| 7R0C | C | arrestin-2–V2R (AVP) | Core-engaged | Yes | 19.24 |
| 7SRS | C | arrestin-2–5-HT2B (LSD) | Core-engaged | Yes | 19.31 |
| 8ZYY | B | arrestin-2–NTSR1 (SBI-553) | Core-engaged | Yes | 19.38 |
| 9ZZO | C | arrestin-2–$\kappa$-opioid receptor (MP1104) | Core-engaged | Yes | 19.41 |
| 9IJS | C | arrestin-2–GPR52 (ligand-free) | Core-engaged | Yes | 19.52 |
| 22JX | C | arrestin-2–mGlu5 | Tail-engaged | Yes | 19.86 |
| 8ZYU | B | arrestin-2–NTSR1 (SBI-553) | Core-engaged | Yes | 20.11 |
| 6TKO | B | arrestin-2–$\beta_1$AR (formoterol) | Core-engaged | Yes | 20.23 |
| 8AS2 | A | arrestin-2 + CCR5 phosphopeptide (4P) | Phosphopeptide | Yes | 21.68 |
| 8AS3 | A | arrestin-2 + CCR5 phosphopeptide (6P) | Phosphopeptide | Yes | 21.97 |

### Supplementary Data 4

To investigate the detachment of the C-tail from its basal binding site, we used a set of complementary CVs designed to capture different aspects of C-tail disengagement, with particular focus on the $\beta 20$ strand. The main OPES Explore bias was applied in all replicas as a single multidimensional bias and included three CVs: a hydrogen-bond-based CV describing formation and disruption of the inter-strand hydrogen-bond network $\mathrm{HB}_{\mathrm{diff}}$, a side-chain packing CV reporting on the compactness of the $\beta 1/\beta 20$ interface $\mathbf{c}_{\mathrm{sidechain}}$ and a $\beta 20$–core coordination/separation CV $\mathbf{d}_{\beta 20\text{-core}}$. Also, water-coordination CVs were applied separately in all replicas to enhance sampling of local solvent exposure at the $\beta 1$ interface. Additional water-coordination CVs were introduced progressively across replicas to improve the sampling of selected local interactions during $\beta 20$ displacement. The OPES Explore barrier for the main multidimensional bias was set to 75 kJ/mol. The water-coordination CVs included in all replicas used a barrier of 4 kJ/mol, whereas the additional water-coordination CVs introduced progressively across replicas used a barrier of 3 kJ/mol.

**Table S3:** Biasing scheme used for the C-tail detachment simulations. All replicas were biased using the main collective variables $\mathrm{HB}_{\mathrm{diff}}$, $\mathbf{c}_{\mathrm{sidechain}}$, and $\mathbf{d}_{\beta 20\text{-core}}$. Water-coordination CVs describing hydration of K10, K11, and A12 were included in all replicas. The additional variables listed in the table are water-coordination CVs that differ between replicas. Here, $w_{\mathrm{X\_Y}}$ denotes the coordination of water oxygen to atom Y of residue X. The column labelled "Replica temperature" indicates the highest temperature reached during each replica, starting from the thermostat temperature of 300 K.

| Replica | Main CVs | Hydration CVs | Additional hydration CVs | Replica temperature |
|---|---|---|---|---|
| 0 | $\mathrm{HB}_{\mathrm{diff}}$, $\mathbf{c}_{\mathrm{sidechain}}$, $\mathbf{d}_{\beta 20\text{-core}}$ | $w_{\mathrm{K10\_C}}$, $w_{\mathrm{K11\_O}}$, $w_{\mathrm{A12\_C}}$ | – | – |
| 1 | $\mathrm{HB}_{\mathrm{diff}}$, $\mathbf{c}_{\mathrm{sidechain}}$, $\mathbf{d}_{\beta 20\text{-core}}$ | $w_{\mathrm{K10\_C}}$, $w_{\mathrm{K11\_O}}$, $w_{\mathrm{A12\_C}}$ | $w_{\mathrm{I386\_CD1}}$ | 305 K |
| 2 | $\mathrm{HB}_{\mathrm{diff}}$, $\mathbf{c}_{\mathrm{sidechain}}$, $\mathbf{d}_{\beta 20\text{-core}}$ | $w_{\mathrm{K10\_C}}$, $w_{\mathrm{K11\_O}}$, $w_{\mathrm{A12\_C}}$ | $w_{\mathrm{I386\_CD1}}$, $w_{\mathrm{F391\_CE1}}$ | 310 K |
| 3 | $\mathrm{HB}_{\mathrm{diff}}$, $\mathbf{c}_{\mathrm{sidechain}}$, $\mathbf{d}_{\beta 20\text{-core}}$ | $w_{\mathrm{K10\_C}}$, $w_{\mathrm{K11\_O}}$, $w_{\mathrm{A12\_C}}$ | $w_{\mathrm{I386\_CD1}}$, $w_{\mathrm{F391\_CE1}}$, $w_{\mathrm{F391\_CD1}}$ | 315 K |
| 4 | $\mathrm{HB}_{\mathrm{diff}}$, $\mathbf{c}_{\mathrm{sidechain}}$, $\mathbf{d}_{\beta 20\text{-core}}$ | $w_{\mathrm{K10\_C}}$, $w_{\mathrm{K11\_O}}$, $w_{\mathrm{A12\_C}}$ | $w_{\mathrm{I386\_CD1}}$, $w_{\mathrm{F391\_CE1}}$, $w_{\mathrm{F391\_CD1}}$ $w_{\mathrm{F391\_CZ}}$ | 320 K |
| 5 | $\mathrm{HB}_{\mathrm{diff}}$, $\mathbf{c}_{\mathrm{sidechain}}$, $\mathbf{d}_{\beta 20\text{-core}}$ | $w_{\mathrm{K10\_C}}$, $w_{\mathrm{K11\_O}}$, $w_{\mathrm{A12\_C}}$ | $w_{\mathrm{I386\_CD1}}$, $w_{\mathrm{F391\_CE1}}$, $w_{\mathrm{F391\_CD1}}$ $w_{\mathrm{F391\_CZ}}$, $w_{\mathrm{V8\_O}}$ | 325 K |
| 6 | $\mathrm{HB}_{\mathrm{diff}}$, $\mathbf{c}_{\mathrm{sidechain}}$, $\mathbf{d}_{\beta 20\text{-core}}$ | $w_{\mathrm{K10\_C}}$, $w_{\mathrm{K11\_O}}$, $w_{\mathrm{A12\_C}}$ | $w_{\mathrm{I386\_CD1}}$, $w_{\mathrm{F391\_CE1}}$, $w_{\mathrm{F391\_CD1}}$ $w_{\mathrm{F391\_CZ}}$, $w_{\mathrm{V8\_O}}$, $w_{\mathrm{V387\_O}}$ | 330 K |
| 7 | $\mathrm{HB}_{\mathrm{diff}}$, $\mathbf{c}_{\mathrm{sidechain}}$, $\mathbf{d}_{\beta 20\text{-core}}$ | $w_{\mathrm{K10\_C}}$, $w_{\mathrm{K11\_O}}$, $w_{\mathrm{A12\_C}}$ | $w_{\mathrm{I386\_CD1}}$, $w_{\mathrm{F391\_CE1}}$, $w_{\mathrm{F391\_CD1}}$ $w_{\mathrm{F391\_CZ}}$, $w_{\mathrm{V8\_O}}$, $w_{\mathrm{V387\_O}}$, $w_{\mathrm{E389\_O}}$ | 335 K |

To assess whether the main detachment CVs were efficiently sampled, we monitored their time evolution in the C-tail detachment OneOPES simulations. The analyzed CVs correspond to the three components of the main multidimensional OPES Explore bias: a hydrogen-bond-based CV describing formation and disruption of the inter-strand hydrogen-bond network $\mathrm{HB}_{\mathrm{diff}}$, a side-chain packing CV reporting on the compactness of the $\beta 1/\beta 20$ interface $\mathbf{c}_{\mathrm{sidechain}}$ and a $\beta 20$–core coordination/separation CV $\mathbf{d}_{\beta 20\text{-core}}$. Their broad sampling over time indicates that the simulations explored both compact and displaced $\beta 20$ configurations.

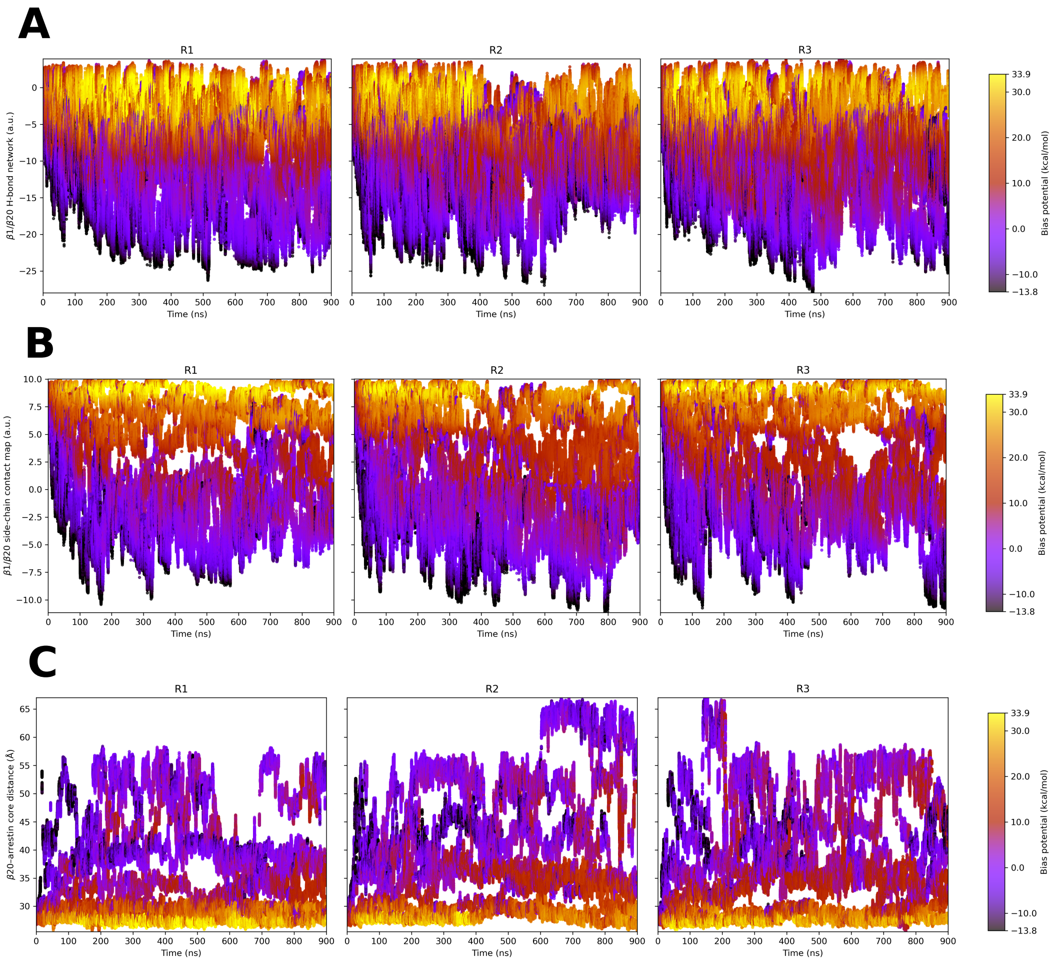


**Figure S4: Sampling of the main CVs in the C-tail detachment OneOPES simulations. (A)** Sampling of the hydrogen-bond–based CV describing formation and disruption of interstrand hydrogen-bond network as a function of time. **(B)** Sampling of the side-chain packing CV reporting on the compactness of the $\beta 1/\beta 20$ interface as a function of time. **(C)** Sampling of the structural separation CV defined as a distance between the geometric center of $\beta 20$ and the geometric center of C$\alpha$ atoms forming $\beta$ strands 1–19 of protein as a function of time. For each panel data points are shown for replica 0 and colored according to the accumulated bias potential.

In addition, we monitored two projection-based descriptors to follow $\beta 20$ displacement along the sampled detachment pathway: axial displacement from the attached-state position along the detachment direction (`pp.proj`) and lateral displacement away from this direction (`pp.ext`). Although the axial projection coordinate was not part of the main multidimensional bias, it is closely related to the biased $\beta 20$–arrestin core separation CV and was used as the projection coordinate for the reweighted free-energy profile. We therefore explicitly inspected its sampling before projecting the reweighted trajectories onto this coordinate.

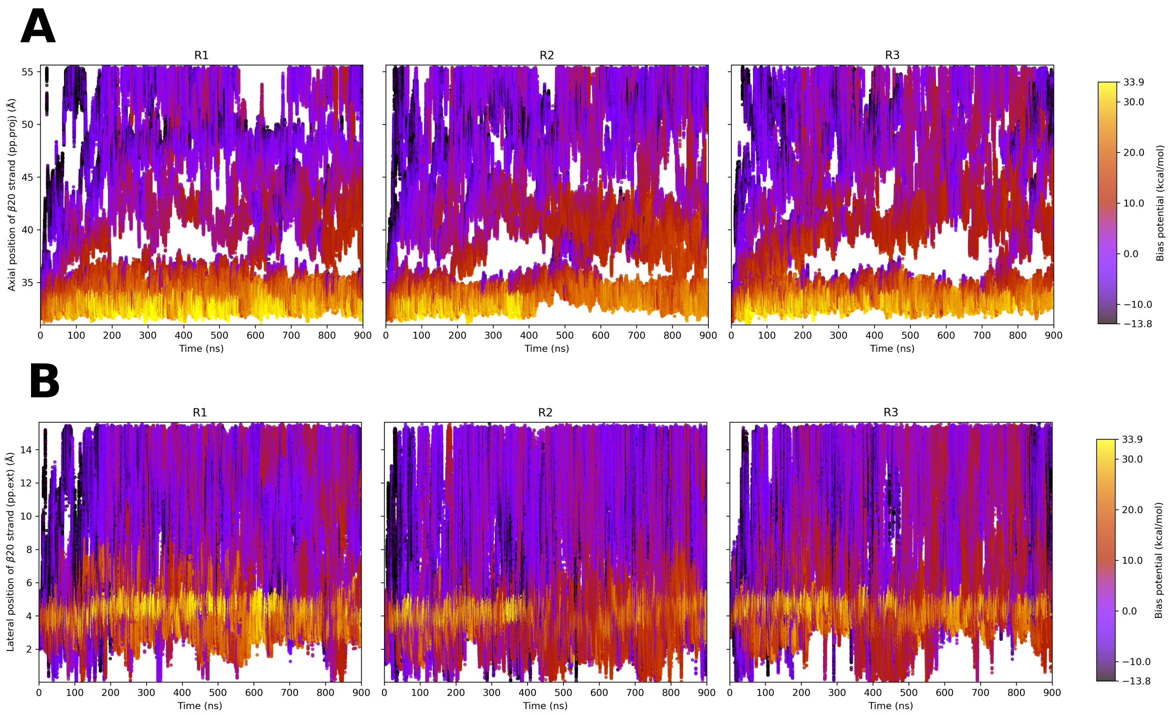


**Figure S5: Sampling of the CVs used for the description of funnel (cylindrical) restraints in the C-tail detachment OneOPES simulations. (A)** Sampling of the axial position of $\beta 20$ strand (pp.proj) as a function of time. **(B)** Sampling of lateral position of $\beta 20$ strand (pp.ext) as a function of time. For both panels data points are shown for replica 0 and colored according to the accumulated bias potential.

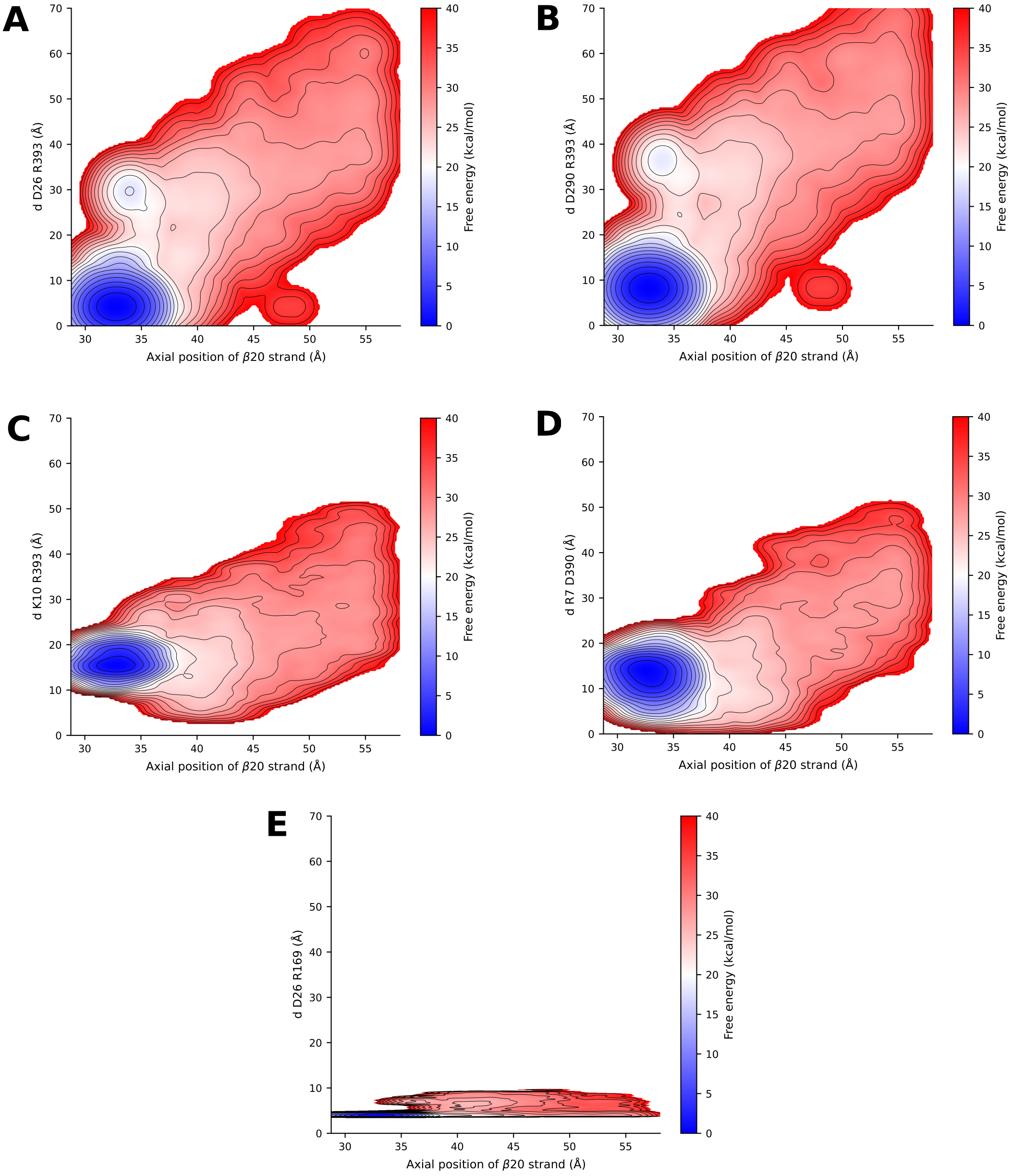


**Figure S6: 2D FES monitoring the axial position of $\beta$20 strand (pp.proj) and selected intramolecular contact distances.** **(A)** D26–R393 **(B)** D290–R393 **(C)** K10–R393 **(D)** R7–D390 **(E)** D26–R169. Low pp.proj values correspond to the attached $\beta$20 state, whereas higher pp.proj values correspond to progressive $\beta$20 displacement from the protein core. The attached state coincides with compact basal contacts, while $\beta$20 displacement is associated with increased contact distances and disruption of the autoinhibited interaction network. The global free-energy minimum is located in the attached state.
